# Left or right: Handedness in free-ranging Hanuman langurs (*Semnopithecus entellus*) residing in an urban ecosystem

**DOI:** 10.1101/2023.03.25.534213

**Authors:** Akash Dutta, Dishari Dasgupta, Arnab Banerjee, Sk Anzar Hasnain, Debadrita Sen, Milan Sahadevan Kuleri, Pritha Bhattacharjee, Manabi Paul

**Affiliations:** Department of Environmental Science, University of Calcutta, Kolkata, India; Department of Biological Sciences, Indian Institute of Science Education and Research Kolkata, India; Agricultural and Ecological Research Unit, Biological Science Division, Indian Statistical Institute, Kolkata, India; Department of Zoology, Dinabandhu Andrews College, Kolkata, India; Department of Biological Sciences, Indian Institute of Science Education and Research, Berhampur, India; Department of Environmental Science, Ashutosh College, Kolkata, India

**Keywords:** Handedness, Free-ranging, Hanuman langur, Postural Origin Theory, Urban ecosystem

## Abstract

Examining manual lateralization (handedness) in nonhuman primates might be an interesting approach to gaining insight into the evolution of asymmetry in humans. Moreover, handedness could also reflect the effect of environmental alterations on the free-ranging animals who are forced to live with anthropogenic interferences. Despite addressing the handedness among monkeys and apes, only a few studies have focused on these free-ranging urban-adapted nonhuman primates, which could challenge our perception of habitat loss and deforestation. Here, we conducted 193 field-based experimental trials with two experimental tasks, one unimanual (simple reaching) and one bimanual (tube task) to explore manual lateralization in a highly human-provisioned group of free-ranging Hanuman langur (*Semnopithecus entellus*). Experimental outcomes revealed an asymmetrical hand-use distribution, with a bias toward the left hand. As bimanual tasks evoked a higher degree of lateralization, these tasks seem to be more suited to study manual laterality, and our results also highlight the significance of experimental tasks in establishing hand preference in langurs. Furthermore, this study also reveals that such lateralization developed with age as adults distinctly displayed their preference toward left-hand usage in contrast to juveniles and subadults who used both hands comparably. Mostly considered to be arboreal, the langurs of our study group spend a considerable amount of time with humans on the ground, thereby portraying a terrestrial tendency. Postural Origin Theory states that terrestrial animals tend to use their right hand and arboreal their left. Therefore, here the presence of group-level left-hand biasness in the adult langurs of Dakshineswar creates a dilemma in the Postural Origin Theory.

## Introduction

Behavioural asymmetry due to brain lateralization such as handedness (manual laterality) has been of great interest among researchers across the globe (Zhao et.al.,2010). It was believed that such asymmetry associated with right-handedness was unique to human beings as 90% of adult humans preferred to use their right hand for several tasks (Annet, 2002; Corballis, 2002; Crow,2004; Cashmore, 2009;). Such right-handed bias corresponds to the specialization of the cerebral hemisphere – the left hemisphere, which controls motor actions required for the completion of a given task. However, the drive behind the evolution of brain lateralization remains unclear, for which, various non-human animal models have been targeted to study their task-oriented cognitive abilities (Ghirlanda & Vallortigara 2004). And there is a growing corpus of evidence suggesting that such asymmetry as a result of brain lateralization is present in other vertebrates as well (Rogers 2002; Rogers & Andrew 2002; Vallortigara & Rogers 2005; Rogers,2009). Most of these studies focus on the hand preference of our closest relatives, non-human primates (NHP) using non-invasive methods like long-term observation and unimanual and/or bimanual tasks to understand the origin and evolution of brain asymmetry. Unimanual tasks (also known as simple reaching tasks) require the involvement of one forelimb, in contrast to bimanual tasks where both the forelimbs remain involved for the completion of a given task (eg. Tube task) (Hopkins & Morris, 1993).

W.D. Hopkins carried out a long-term study on captive chimpanzees (*Pan troglodytes*) where he observed the handedness of the species with the help of a ‘tube task’. In the task, chimpanzees needed to hold a PVC tube in one hand and remove the smeared peanut butter from the inside of the tube using the other hand (dominant hand). A population-level right-hand biasness was found with adults being more lateralized than adolescents and juveniles (Hopkins, 1995). Similar studies on gorillas, bonobos, and ring-tailed lemurs reported population-level right-hand biasness. Gorillas showed a right-hand biasness for their intraspecific gestures as well as for food-acquiring tasks (Byrne & Byrne,1991; Prieur et.al.,2016), bonobos showed a right-hand biasness for fulfilling tasks requiring tool usage (Bardo et.al.,2015), and in ring-tailed lemurs, a group-level right-hand biasness was observed for both unimanual and bimanual tasks (Regaiolli et.al.,2016).

However, a study was done on the wild arboreal old-world monkey, Sichuan snub-nosed monkey (*Rhinopithecus roxellana*) revealed a population-level left-hand biasness for the bimanual tube task (Zhao et.al.,2012). Similar population-level left-hand biasness was also found in deBrazza’s monkey (*Cercopithecus neglectus*), and Orangutans (*Pongo pygmaeus*), suggesting that terrestrial primates tend to prefer right-hand and arboreal primates left-hand for bimanual tasks (Schweitzer et al., 2007; Hopkins et.al.,2003). These findings are in concordance with the ‘Postural Origin Theory’ which states that arboreal primates mostly used their right hand to support themselves and their left hand to grab, snatch, and reach objects. In the course of evolution, as the primates gained an upright posture to become more terrestrial, they started using their right hand for tool-usage instead of providing support to the body. This change in posture of the primates from being arboreal to being terrestrial played a pivotal role in the brain lateralization found in humans (MacNeilage et.al.,1987, MacNeilage,1998).

Besides bimanual tasks, simple reaching tasks are also common for studying handedness in NHP models. However, according to the ‘task-complexity theory’, simple reaching tasks often do not elicit a directional handedness at the population level (Fagot & Vauclair,1991). It posits that the difficulty level of the task significantly affects population-level hand biasness. Complex tasks require the subjects to do cognitively demanding activities in contrast to simple reaching tasks (Fagot & Vauclair,1991). However, the outcomes remained inconsistent for individual and population levels and varied largely for different species of NHP (Meguerditchian et al.,2013).

Even though several studies were done on NHP species belonging to the catarrhine lineage, only a single study has been done on captive Hanuman langurs. Here, langurs were considered as terrestrial NHP and showed a left-hand biasness for the tube task, thereby disagreed with the Posture Origin Theory (Caspar et. al.,2022). However, similar studies on terrestrial rhesus macaques revealed a right-hand biasness, and agreed with the Posture Origin Theory (Westergaard and Soumi, 1996). Interestingly, a conundrum lies regarding the substrate use of *Semnopithecus entellus*, while most of the studies have considered langurs to be arboreal (Karanth et.al., 2017; Adhikari & Dhakal, 2018; Ahamed & Dharmaretnam, 2003; Bhattacharyya et. al., 2008; Egi et.al., 2007; Kashyap et.al., 2011; Narasimmarajan et.al., 2012; Patil & Modse, 2018; Rahman et.al., 2015; Sayers & Norconk, 2008; Sushma & Singh, 2006), few have cited them as semi-terrestrial (B.A. Patel,2010; Rahman et.al.,2015) and terrestrial (Caspar et.al.,2022). Now, according to Posture Origin Theory, langurs are expected to show a left-hand biasness if they are arboreal and a right-hand biasness if terrestrial (MacNeilage et.al.,1987, 1998). Our previous study (Dasgupta et.al.,2021) revealed that the langur troop in Dakshineswar (DG), West Bengal spends a considerable amount of time with humans, and largely thrive on human provisioning, thereby have developed a keen interest in human processed food items. It is of high expectation that a terrestrial species could interact more with humans in contrast to an arboreal species. Therefore, we were curious to explore the hand usage of such a human-provisioned langur troop who have a tendency to be terrestrial. Here, in this study, free-ranging Hanuman langurs (*Semnopithecus entellus*) of DG have been used as the study subject to perform a field-based experimental task (modified version of the tube task) to explore the handedness at the population level. Moreover, this study highlights the correlation between manual laterality and age groups of DG langurs.

## Methodology

### Study area and Subject

Free-ranging gray langur (*Semnopithecus entellus*) groups were identified through regular census between August 2020 and November 2021 in different parts of West Bengal, India, of which three distinct langur groups (one in Dakshineswar (22.655721, 88.357741), one in Nangi (22.507134, 88.211681), and one in Kurumba, Birbhum (23.720748, 87.777782) were selected for long-term observations, considering various level of human interferences received by these langurs. Among these three locations, Dakshineswar has been considered as an urban landscape having ~ 77 percentage of built-up area in contrast to Nangi and Kurumba (built up area ~ 23.31 % and ~ 10.63% respectively) (**Figure 1**). Besides, langur troop in Dakshineswar (DG) receive notable human attention because of their deity values, and are frequently observed to extend their hands to receive human-offered food items (Dasgupta et al. 2021; Dasgupta et al., manuscript in preparation).

**Figure 1:**
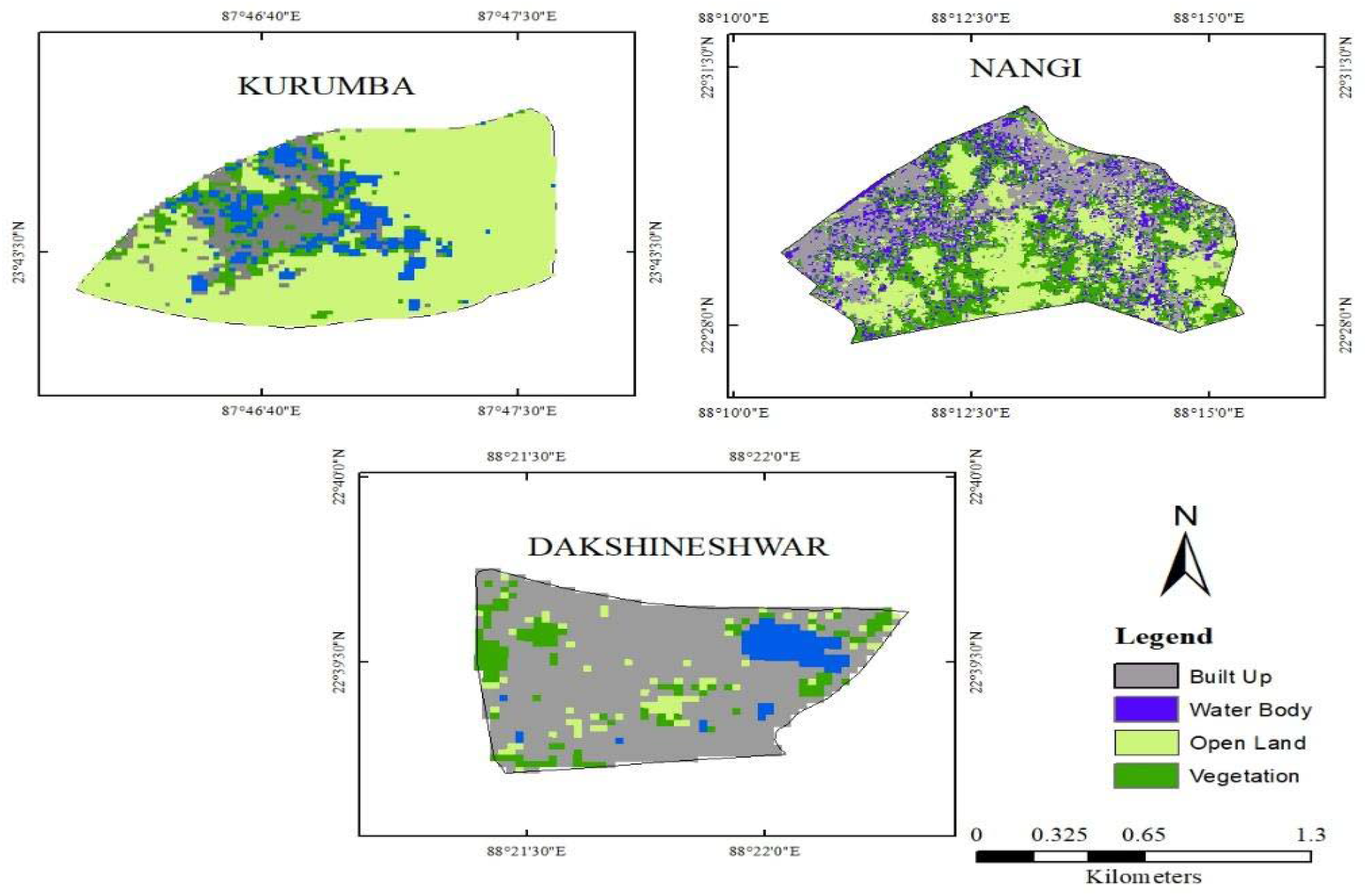
Supervised classification map showing vegetation, open land, waterbody, and built-up area of three study sites, Kurumba, Nangi, and Dakshineshwar. Here, Dakshineswar has 77 percentage of built-up area in contrast to Nangi and Kurumba (built up area ~ 23.31 % and ~ 10.63% respectively).

Hence for this study we have selected DG to explore handedness traits within an urban ecosystem. DG consists of total 35 individuals (1 adult male, 12 adult females, 6 Subadults, 8 juveniles, and 8 infants) of which all adults, sub-adults, and juveniles have been chosen randomly for the commencement of these field-based experiments. We have excluded the infants from the experiment as they rarely interact with humans and remained dependent on adult females for their feeding and movements. Therefore, it becomes difficult to involve them without intimidating the adult langur in the experiment. Total 193 field-based trials have been recorded between August 2021 to July 2022, of which only 156 trials have been considered as successful events for the final analysis.

### Experiment design

We have designed a field-based experiment to understand the hand usage in langurs of highly human-provisioned DG that requires the involvement of hands for the completion of given tasks. For each experiment, we have used a transparent bottle with a height of 18 cm and a diameter of 22 cm, which has been baited with a food item. As buns are easily available and one of the preferred food items of these DG langurs (Dasgupta et al., 2021), here, we have used them as bait and kept inside the bottle in the absence of focal langurs (**Figure 2, Panel I**). For each trial, the observers position themselves with the bottle in front of the subject (individual langur, randomly selected for the experiment), so that the bottle is at equidistance from both the hands of the subject (**Figure 2**, **Panel II**, ESM Video clip).

**Figure 2:**
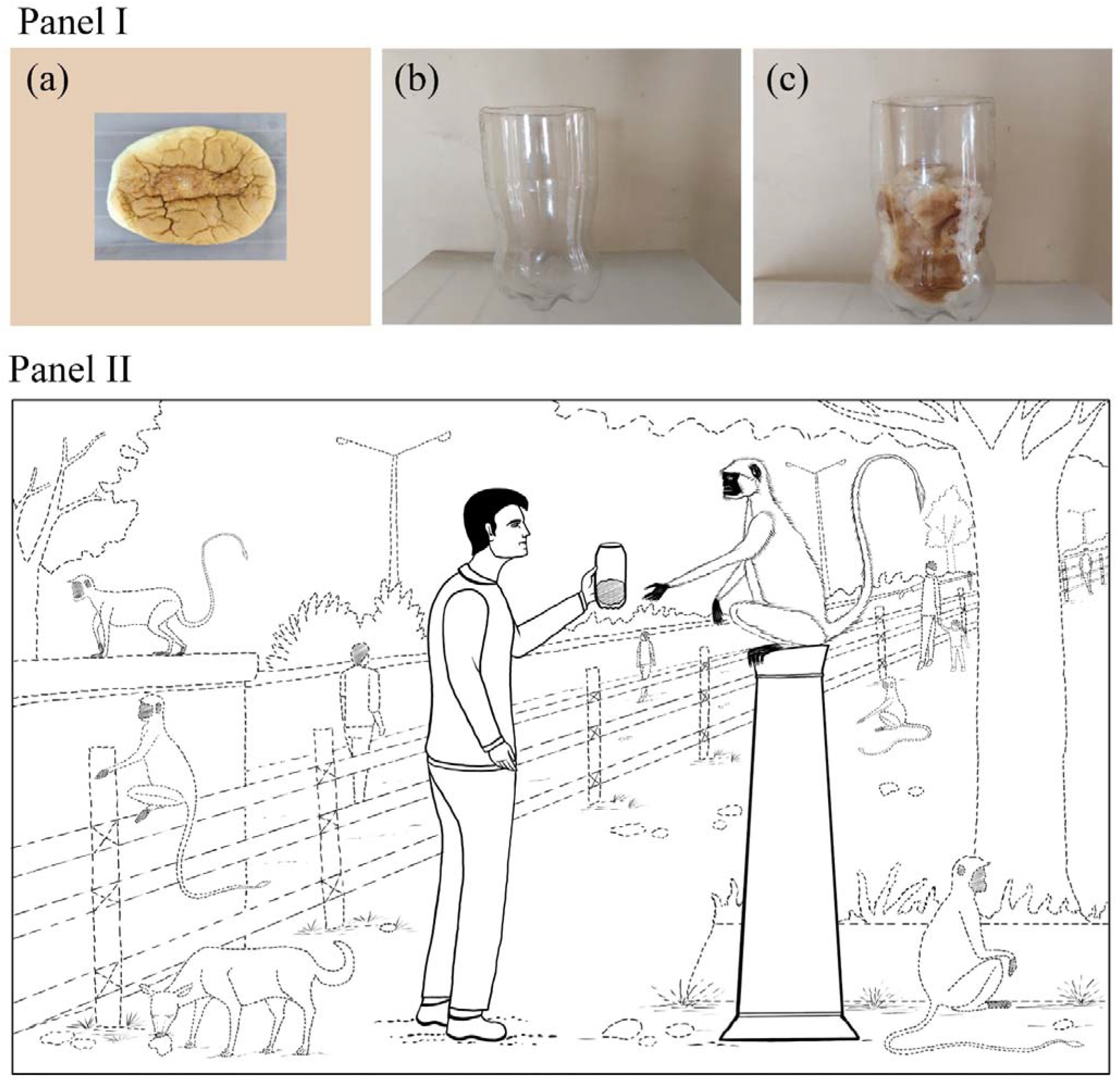
Panel I represents the experiment setup, (a) bun, (b) a transparent bottle with a height of 18 cm and a diameter of 22 cm, and (c) complete setup for the experiments. Panel II represents a 2D visualization of the task-based hand-choice field experiment.

The experiment is largely divided into two tasks: i) T1: a unimanual simple reaching task that requires the langur to collect the bottle from the experimenter and ii) T2: A bimanual task where the langur has to take out the bait from the bottle. We have recorded the ‘Start time’ (ST) as soon as the subject has taken the bottle from the observer and went on until the subject completely removed the bait from the bottle. While performing the second task, often the bait has been removed partially in smaller pieces and consumed by the subject simultaneously. The experimenter has recorded the ‘End time’ (ET) only when the entire bait has been removed from the given bottle and has been denoted as the completion of the second task. All the trials are recorded in Sony Digital HD Video Camera Recorder (HDR-CX470) with 60x clear image zoom which have been conducted either in the morning (0900-1200) or in the evening (1500-1800) session.

### Data Decoding

All the videos have been decoded, noting the usage of body parts (BP) like limbs (hands and legs) and mouth (M) to perform both T1 and T2. Since T2 involves usage of both limbs and mouth for multiple times (to remove the bait completely from the bottle) in contrast to T1 which is a simple reaching task, the observer has also recorded the frequency of using BP separately by means of slow motion and still-frame video analysis. Moreover, hand usages (H) are again sorted into three categories: 1. by using left hand (LH), 2. by using right hand (RH), and 3. by using both hands (BH). For each trial, the task completion time (TC) (latency between ST and ET) is also calculated separately. All the extractive responses, including the unsuccessful attempts at food retrieval, have been grouped into three categories based on the age class of langurs; adults, subadults and juveniles.

### Data Analysis

Manual laterality (handedness) in DG langurs is evaluated based on hand preference and performance (Spinozzi et al., 2007). While hand preference describes the frequency with which the right or left hand is used to extract food, the task completion time (TC) is used to evaluate the performance separately for the right and left hands. We have used six different analyses to assess the manual laterality in these langurs. First, we ran a correlation analysis between the task completion times (TC) and use of hands (H) (including a detailed involvement of right (RH), left (LH), and both hands (BH)), mouth (M) and leg (L) for various life stages of langur. Then we did a principal component analysis (PCA) to determine the importance of these body parts (BP) selection for the completion of given tasks, followed by a t-test analysis and Jaccard similarity index. While the t-test compared the proportion of left and right hands used per subject, the JS index checked the difference between the frequency of right and left-hand usage. Jaccard Similarity Index is a measure of similarity for the two sets of data, with a range from zero (0) to one (1). Values closer to 1, suggested increased similarity between the two sets and the Jaccard distance is complement of the Jaccard index, which is a metric for measuring the dissimilarity of two sets. By deducting the Jaccard Index from 1, Jaccard distance can be obtained.

#### Handedness index

We also measured their directional handedness index (HI). Here, we used the formula (R-L)/(R+L) to calculate HI for each individual langur, where R and L represent the total number of right- and left-hand extractive responses, respectively (Spinozzi et al., 2007). HI ranges between +1.0 (completely right-handed) to −1.0 (completely left-handed), thereby revealing the preference for manual direction. The strength of hand preference is assessed by the absolute value of the HI (ABS-HI), which is direction-independent. The stronger the laterality, the closer the ABS-HI is to 1. A univariate analysis followed by a post-hoc Tuckey test was used to interpret these ABS-HI scores.

Finally, two sets of linear models were used to determine the relationship between task completion (TC) and BP use. Both models used the TC as the response variable. While we incorporated hand (H), mouth (M), and leg (L) as the fixed variables in the first model, the second model used the right hand (RH), left hand (LH), and both hands (BH) as the fixed effects. All the statistical tests were carried out using R (version 4.2.2) (R Core Team 2022).

### Ethical Note

No gray langurs were harmed during this work. All work reported here was purely observation-based and did not involve direct handling of gray langurs in any manner, therefore, was in accordance with approved guidelines of animal rights regulations of the Government of India. The research reported in this paper was sanctioned by DST-INSPIRE, Government of India (approval number: DST/INSPIRE/04/2018/001287, dated 24th July 2018). Due permission was taken from the Principal Chief Conservator of Forests (PCCF), West Bengal, India (869/WL/4R-43/2021).

## Results

Correlation plot 1 (**Figure 3a**) reveals that TC has the highest association with hand (H) use (*r* = 0.816, *p* < 0.01), while plot 2 (**Figure 3b**) represents left hand (LH) as the most preferred forelimb (dominant hand) for the task completion (*r* = 0.82, *p* < 0.01). A scree plot and a biplot were produced from the PCA (**Figure 4**). Scree plot (**Figure 4a**) shows that the first two components (PC1 and PC2) can effectively represent 90.82% of all the observations while the biplot (**Figure 4b**) reveals that the hand (H) is most important along PC1 (loading = 0. 81) followed by left hand (LH) (loading = 0.38). Moreover, a t-test analysis, comparing the proportion of left and right hand used per subject revealed a significant left-hand use (*t* = 3.78, *p* < 0.001) (**Figure 5**). The Jaccard Similarity Index is 0.04, having a Jaccard Distance of 0.9.

**Figure 3:**
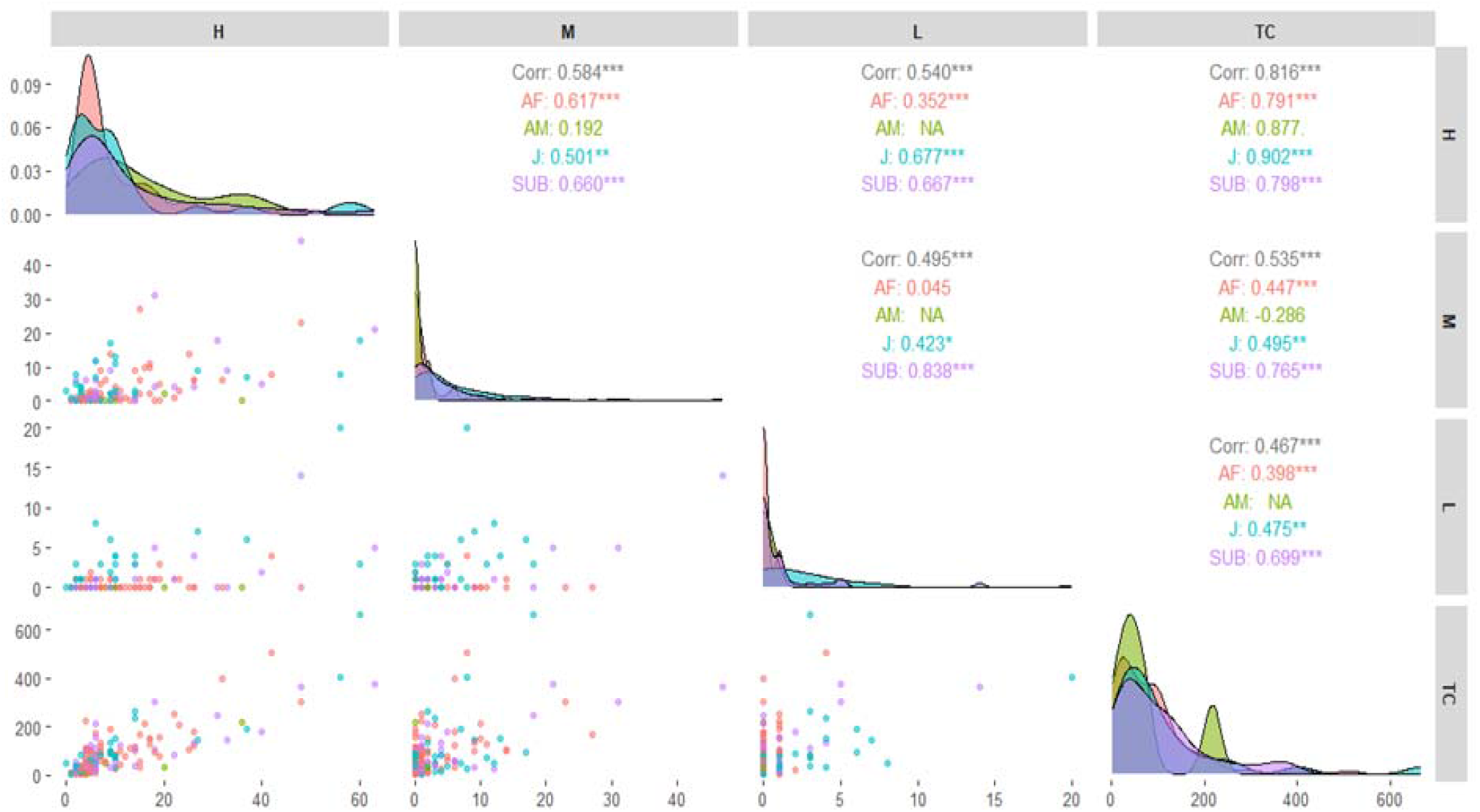
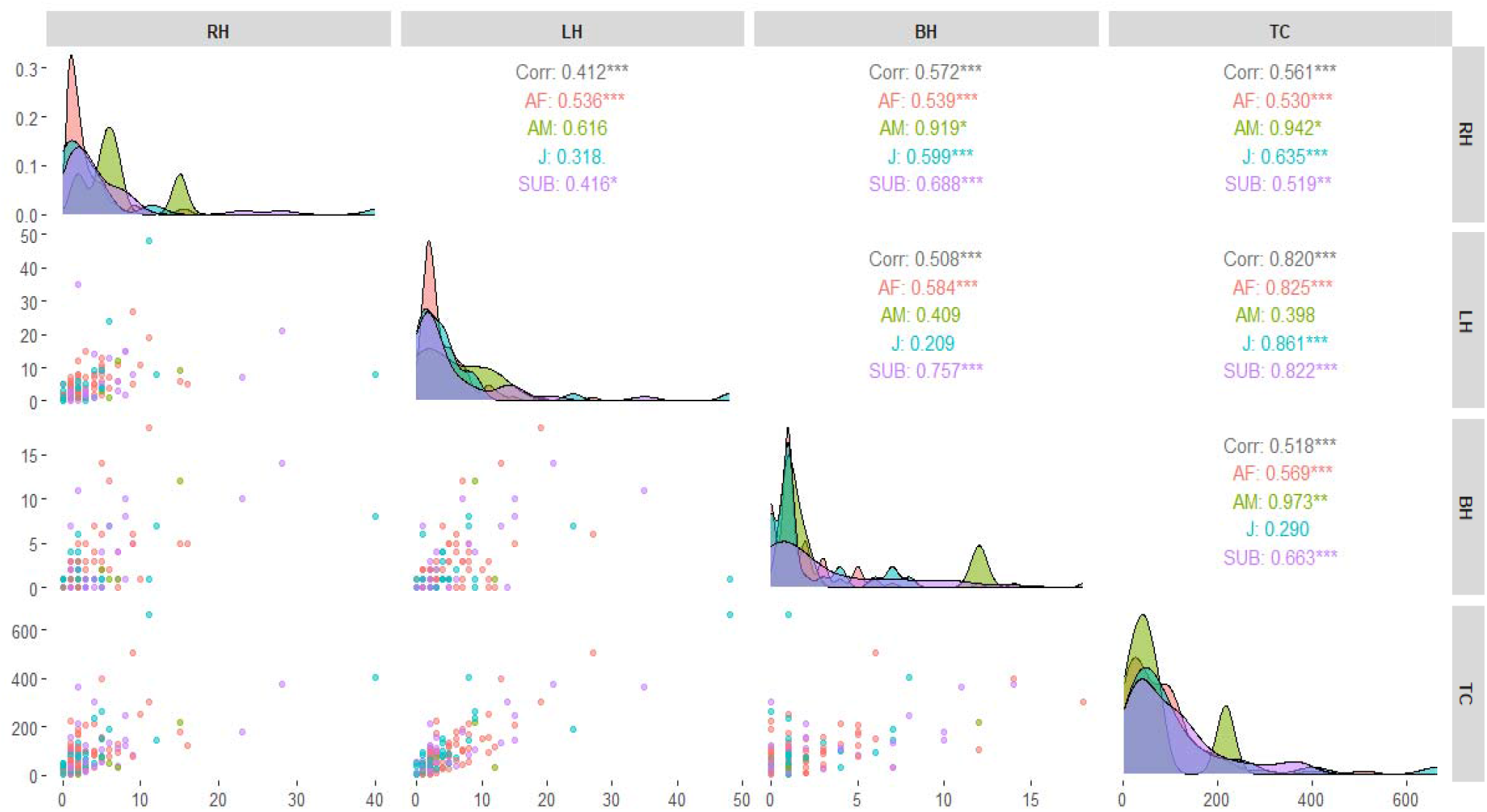
(a) Correlation plot of hand(H), mouth (M), and leg (L) with task completion time (TC). This Correlation plot reveals that TC has the highest association with hand (H) use (*r* = 0.816, *p* < 0.01). (b) Correlation plot of left hand (LH), right hand (RH), and both hand (BH) with task completion time (TC). This plot represents the left hand (LH) as the most preferred forelimb (dominant hand) for the task completion (*r* = 0.820, *p* < 0.01).

**Figure 4:**
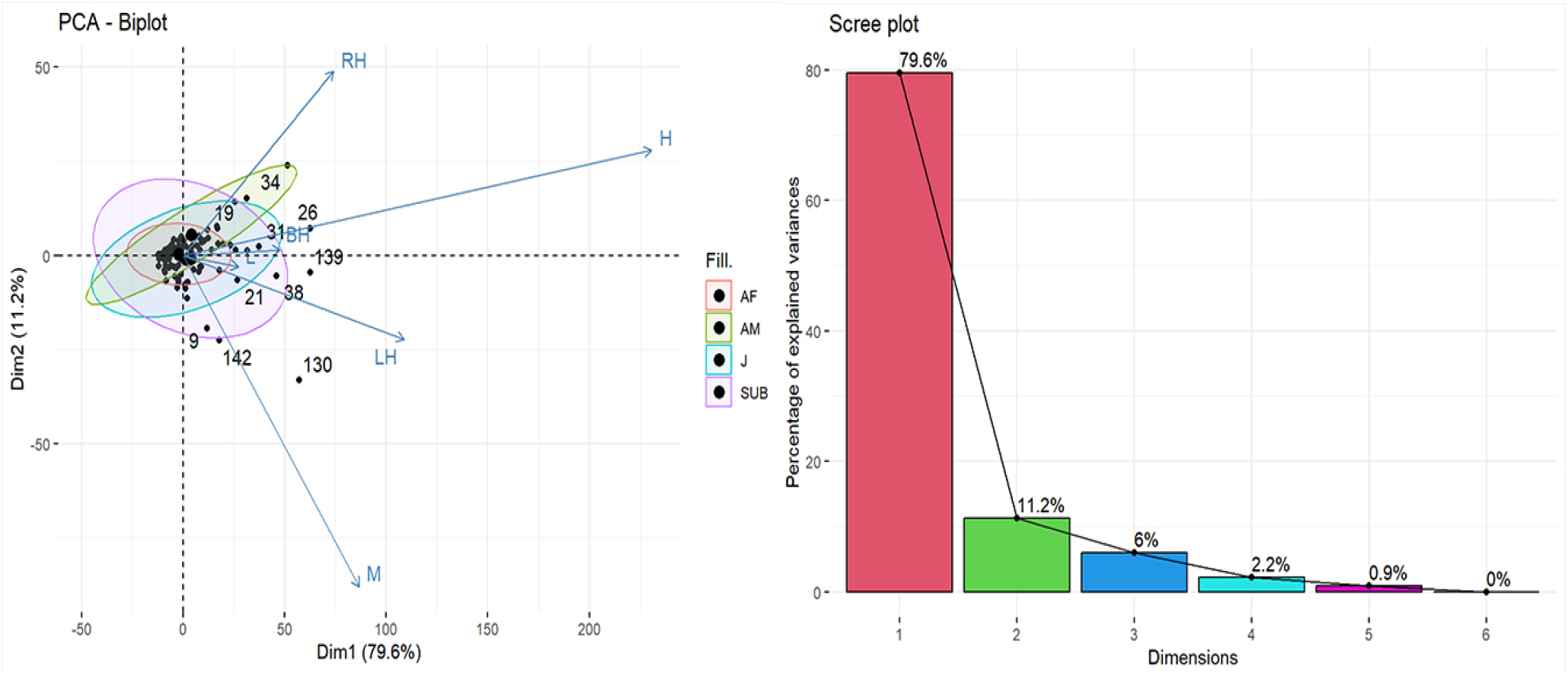
PCA Biplot and Scree Plot. From the scree plot, it can be observed that the first two components (PC1 and PC2) can effectively represent 90.82% of all the observations and the biplot shows that the hand (H) is most important along PC1 (loading = 0. 81) followed by left hand (LH) (loading = 0.38).

**Figure 5:**
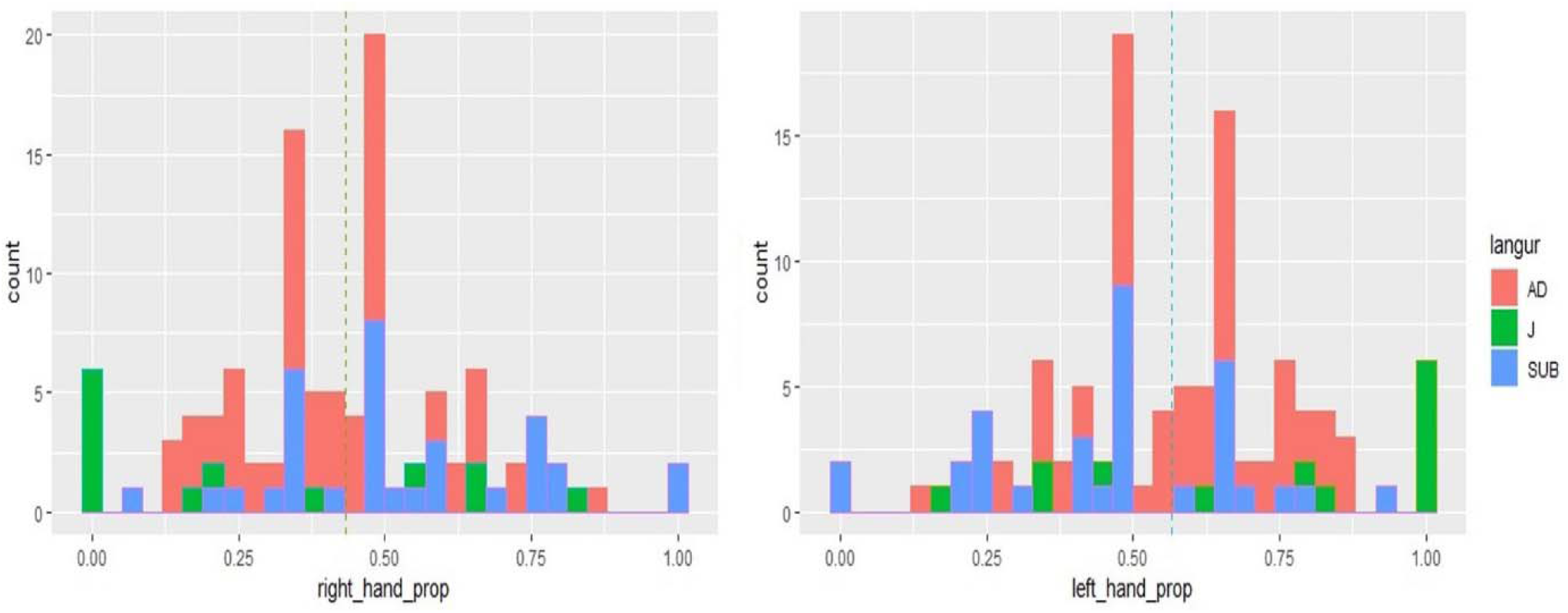
Distribution of proportion use of left and right hand to complete the task. Moreover, a t-test analysis, comparing the proportion of left and right hand used per subject revealed a significant left-hand use (*t* = 3.780,*p* < 0.001)

### Handedness index

A one-sample t-test analysis of the HI score per subject revealed significant hand preference for the age group (mean *HI* = −0.13; *t* = −3.78; *p* < 0.001). While subadult and juvenile failed to show any hand preference, HI shows left handedness in adults (**Figure 6**). Univariate analysis for ABS-HI scores shows significant difference between age group of langurs (*F* = 5.20, *p* < 0.05). The post-hoc analysis (Tuckey Test) revealed significant differences between adult and juvenile (Mean difference = −0.19, *p* < 0.05), as well as between sub-adult and juvenile (Mean difference = −0.19, *p* < 0.05).

**Figure 6:**
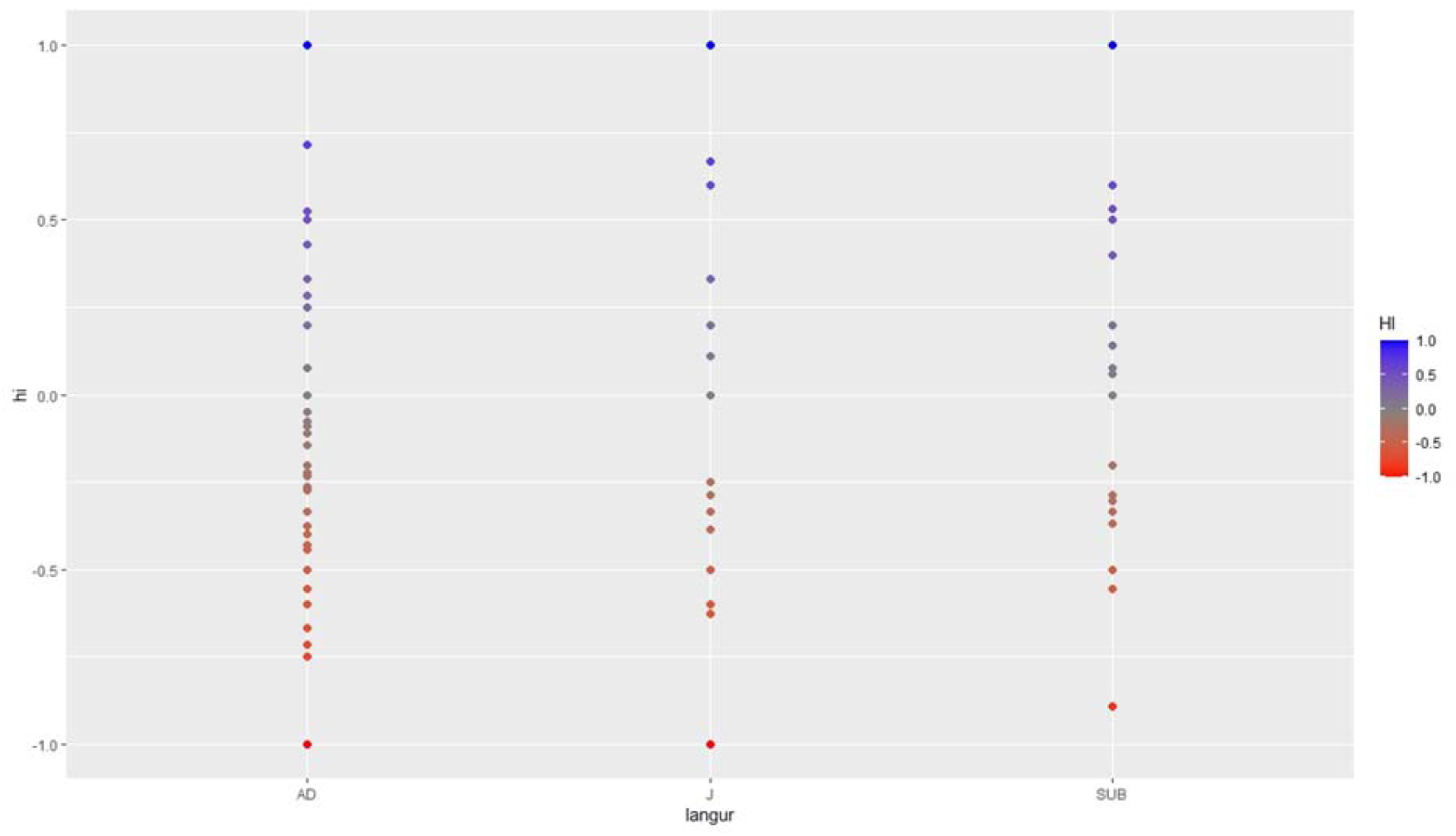
Handedness Index for different age classes of Langur. While subadult and juvenile failed to show any hand preference, HI shows left handedness in adults.

### Linear models

While GLM 1 (model 1) reveals that langurs prefer to use their hand (H) to achieve their goal (parameter estimate = 6.48, *p* < 0.01) (**Table 1a & Figure 7**), GLM 2 (model 2) indicates that the langurs choose their left hand most for task completion as compared to right hand (parameter estimate: *LH* = 11.27 and *RH* = 5.40, *p* < 0.01). The model plot (**Figure 7**) shows the increasing trend of event success with increased use of either appendage. Subplot E shows that LH use has that maximum potential (most steep gradient) leading to event success or task completion (**Table 1b**).

**Table 1a:**
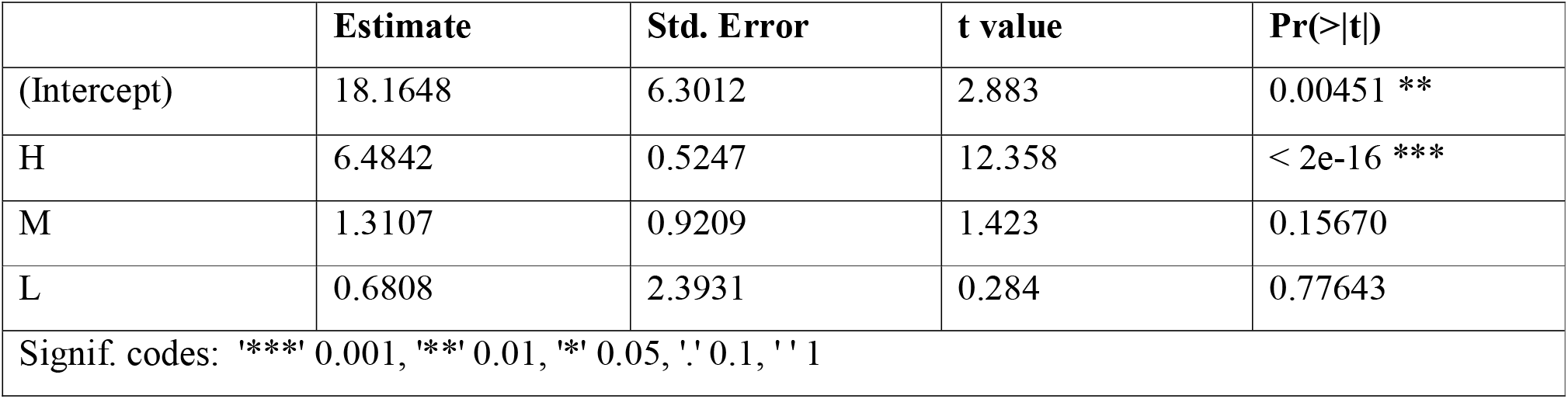
Output of Linear Model 1. LM 1 reveals that langurs prefer to use their hand (H) to achieve their goal (parameter estimate: *H* = 6.48, *p* < 0.01).

**Figure 7:**
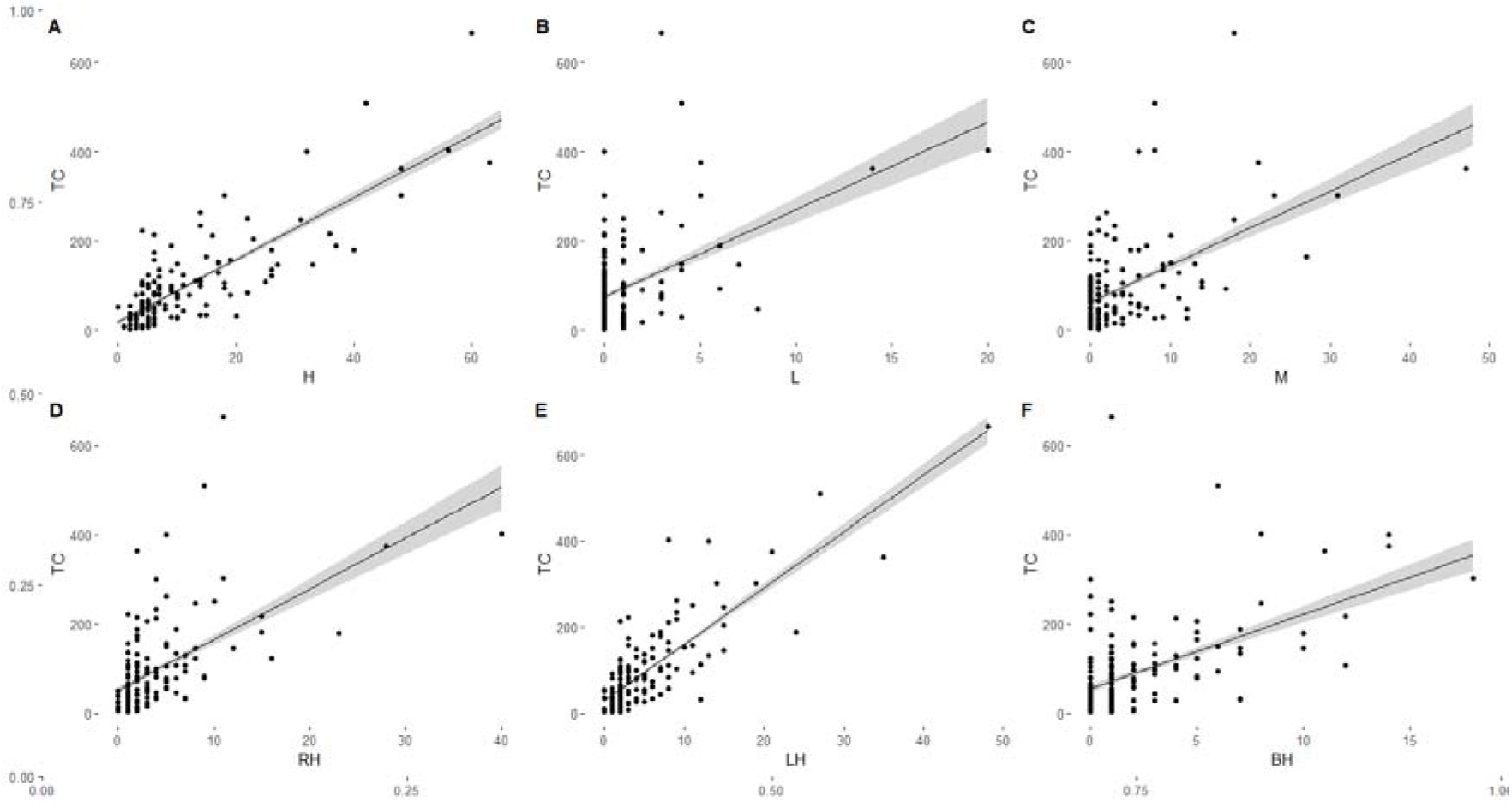
Graphical representation of linear model prediction. While LM 1 reveals that langurs prefer to use their hand (H) to achieve their goal (parameter estimate: *H* = 6.48, *p* < 0.01), LM 2 indicates that the langurs choose their left hand most for task completion as compared to right hand (parameter estimate: *LH* = 11.27 and *RH* = 5.40, *p* < 0.01). The LM plot shows the increasing trend of event success with increased use of either appendage. Subplot E shows that LH use has that maximum potential (most steep gradient) leading to event success or task completion.

**Table 1b:** Linear Model 2. LM 2 indicates that the langurs choose their left hand most for task completion as compared to right hand (parameter estimate: *LH* = 11.27 and *RH* = 5.40, *p* < 0.01).

|  | Estimate | Std. Error | t value | Pr(>\|t\|) |
| --- | --- | --- | --- | --- |
| (Intercept) | 18.3820 | 5.6492 | 3.254 | 0.0014 ** |
| RH | 5.3983 | 1.0499 | 5.142 | 8.27e-07 *** |
| LH | 11.2724 | 0.7878 | 14.308 | < 2e-16 *** |
| BH | 0.1866 | 1.7620 | 0.106 | 0.9158 |
| Signif. codes: '***' 0.001, '**' 0.01, '*' 0.05, '.' 0.1, ' ' 1 |  |  |  |  |

## Discussion

With inconsistent directional handedness at the population level among different species of non-human primates, Hubert et.al., made a pertinent point stating *“Should handedness be considered a dichotomous variable, split between left- and right-handers or between consistent- and inconsistent-handers? Or is it better to conceptualize handedness as continuous rather than categorical, ranging from strongly right to inconsistent to strongly left?”* (Hubert et.al.,2021). Therefore, the primary objective of this study was to determine the handedness of the free-ranging group of Hanuman langurs of Dakshineswar (DG), whether they have a strong right, inconsistent or strong left-hand biasness.

Our 156 successfully executed field-based experimental trials involving a bimanual task revealed a group-level left-handed biasness among DG langurs. The t-tests, HI and linear models reveal that the troop preferred their left hand to complete the given task (**Figure 5, 6, and 7**). This outcome is in agreement with a recent study done on captive, Hanuman langurs where they too reported a left-hand biasness (Casper et.al.,2022). However, a study done on a free-ranging troop of langurs of Ramnagar, Nepal failed to exhibit any such population level handedness (Mittra et.al.,1997). Here, the authors, focussed on activities that required the focal subject to use one hand for its execution like for eating, grooming, reaching etc. However, simple reaching tasks might not always elicit a directional handedness at the population level, as already hypothesized by task-complexity theory (Fagot & Vauclair,1991). Simple reaching task failed to report any population-level biasness for squirrel monkeys as well even though they exhibited left-handedness for catching live fishes (King & Landau, 1993). Therefore, our study was designed in such a way that demanded the focal subject to carry out a complex task in order to extract the reward (bun). Even though some tried to extract the bun by trying to tear open the bottle with their mouth (**Figure 3a**), most of the DG langurs used their left hand to extract the food while holding the bottle with their right hand. As per the Postural Origin Theory (MacNeilage et. al., 1987, MacNeilage,1998), free-ranging langurs, who are considered to be arboreal in their natural habitats (**Ref**), are expected to use their left hand most. The theory states that terrestrial animals prefer to use their right hand most and arboreal their left. But, herein lies the dilemma. These DG langurs, who lost their natural home due to urban sprawling, have learned to survive within human-modified ecosystems. These langurs are largely dependent on human-provisioned food items and considerably interacted with pilgrims (Dasgupta et al. 2021). Owing to their high interactions with humans this troop spends a significant amount of their time on the ground (Dasgupta et al, manuscript in preparation), making this troop more terrestrial rather than arboreal. Therefore, as per the Postural Origin Theory, it is expected that, unlike their natural brethren, DG langurs will show a tendency to use their right hand most. But the group level left-hand biasness here therefore disagrees with the Postural Origin Theory (MacNeilage et.al.,1987, MacNeilage,1998).

Furthermore, the univariate analysis of ABS-HI score reports a significant difference between juvenile, subadult and adult stages of langurs (*F* = 5.199, *p* < 0.05). HI shows left hand biasness among adults but fails to reveal any strong handedness in subadults and juveniles (Figure 5). Altogether, adults seem to be more lateralized for the given experimental tasks than juveniles and subadults, suggesting that hand preference might develop with age. Moreover, subadults tend to use both left and right hand, suggesting that langurs at this life stage tend to be more opportunistic and aim to achieve extra benefit at its best. Previous studies done on other NHPs such as *Callithrix jacchus, R. roxellana* suggests that primate laterality increases with age as the younger population tends to fluctuate between right and left hand for manual actions, showing an inconsistent hand preference in comparison to adults (Hook and Rogers, 2000; Zhao et.al.,2012). However, it is not known exactly at what age handedness develops.

In totality, our study reports that despite the availability of both hands, legs, and mouth, adult langurs of DG notably used their left hand for the successful completion of the task. Therefore, it could be interpreted as an urban adaptation in free-ranging langurs who show a keen interest to be terrestrial for their successful sustenance within urban jungles, but tend to keep their ancestral tendency of using left-hand for the completion of given tasks. Even though our robust study results indicate a clear biasness of handedness, it is essential to replicate this study among more troops of free-ranging Hanuman langurs across different habitats with varying anthropogenic interferences. Moreover, studies should not be restricted to bimanual tasks only, naturally occurring behaviours that require bimanual coordination should also be taken into account in the future. Carrying out a comparative study using the same task among different free-ranging primates would also be of great interest.

## Author Contributions

AD, DD, SAH, DS, and MSK carried out the field work. AD and DD coded the entire data. AD prepared the GIS map. AB supervised and carried out the statistical analyses and AD assisted. AB prepared the correlation matrix, ran PCA and LM & AD prepared the HI to interpret the data. AD, DD and MP conceptualized the study. MP got grants to support the work, designed the fieldwork, and supervised the work. MP, AD, DD, and PB drafted the manuscript.

## Acknowledgments

All the authors acknowledge Mr. Supradipta Dutta for the 2D digital sketch of field experiment and Dr. Anindita Bhadra, Associate Professor, Indian Institute of Science Education and Research-Kolkata, India, for her valuable inputs to the study.

## References

Adhikari, P. P., & Dhakal, P. (2018). Prevalence of gastro-intestinal parasites of Rhesus macaque (Macaca mulatta Zimmermann, 1780) and hanuman langur (Semnopithecus Entellus Dufresne, 1797) in Devghat, Chitwan, Nepal. Journal of Institute of Science and Technology, 22(2), 12–18.

Ahamed, R., & Dharmaretnam, M. (2003). Ranging pattern, feeding and time budget of langurs (Semnopithecus entellus) in a recently established home range at Eastern University Campus, Batticaloa, Sri Lanka. Journal of Science Eastern University of Sri Lanka, 3(1), 1–10.

Ahamed, R., & Dharmaretnam, M. RANGING PATTERN, FEEDING AND TIME BUDGET OF LANGURS (Semnopithecus entellus).

Annett, M. (2002). Handedness and Brain Asymmetry: The Right Shift Theory. Hove: Psychology Press.

Bardo, A., Pouydebat, E., & Meunier, H. (2015). Do bimanual coordination, tool use, and body posture contribute equally to hand preferences in bonobos ?. Journal of Human Evolution, 82, 159–169.

Bennett, A. J., Suomi, S. J., & Hopkins, W. D. (2008). Effects of early adverse experiences on behavioural lateralisation in rhesus monkeys (Macaca mulatta). Laterality, 13(3), 282–292.

Bhattacharyya, T. P., Murmu, A., Chaudhuri, S., & Mazumder, P. C. (2008). Status of four arboreal species of mammals in Darjeeling District, West Bengal, India. Records of the Zoological Survey of India, 108(3), 9–19.

Byrne, R. W., & Byrne, J. M. (1991). Hand preferences in the skilled gathering tasks of mountain gorillas (Gorilla g. berengei). Cortex, 27(4), 521–546.

Cashmore, L. (2009). Can hominin “handedness” be accuartely assessed? Ann. Hum. Biol., 36, 624–641.

Caspar, K. R., Pallasdies, F., Mader, L., Sartorelli, H., & Begall, S. (2022). The evolution and biological correlates of hand preferences in anthropoid primates. Elife, 11, e77875.

Corballis, M.C. (2002). From Hand to Mouth: The Origins of Language. Princeton University Press, Princeton, NJ.

Crow, T. (2004). Directional asymmetry is the key to the origin of modern Homo sapiens (the Broca-Annett axiom): A reply to Rogers‘ review of The Speciation of Modern Homo Sapiens. Laterality, 9, 233–242.

Dasgupta, D., Banerjee, A., Karar, R., Banerjee, D., Mitra, S., Sardar, P.,… & Paul, M. (2021). Altered food habits? Understanding the feeding preference of free-ranging gray langurs within an urban settlement. Frontiers in Psychology, 12, 649027.

Egi, N., Nakatsukasa, M., Kalmykov, N. P., Maschenko, E. N., & Takai, M. (2007). Distal humerus and ulna of Parapresbytis (Colobinae) from the Pliocene of Russia and Mongolia: phylogenetic and ecological implications based on elbow morphology. Anthropological Science, 0705240003–0705240003.

Fagot, J., & Vauclair, J. (1991). Manual laterality in nonhuman primates: a distinction between handedness and manual specialization. Psychological bulletin, 109(1), 76.

Ghirlanda, S., & Vallortigara, G. (2004). The evolution of brain lateralization: a game-theoretical analysis of population structure. Proceedings of the Royal Society of London. Series B: Biological Sciences, 271(1541), 853–857.

He, W., Lu, H., Zhao, K., Song, D., Gai, X., & Gao, F. (2012). Complete genome sequence of a coxsackievirus B3 isolated from a Sichuan snub-nosed monkey.

Hook, M. A., & Rogers, L. J. (2000). Development of hand preferences in marmosets (Callithrix jacchus) and effects of aging. Journal of Comparative Psychology, 114(3), 263.

Hopkins, W. D. (1995). Hand preferences in juvenile chimpanzees: Continuity in development. Developmental Psychology, 31, 619–625.

Hopkins, W. D., & Morris, R. D. (1993). Handedness in great apes: A review of findings. International Journal of Primatology, 14, 1–25.

Hopkins, W. D., Stoinski, T. S., Lukas, K. E., Ross, S. R., & Wesley, M. J. (2003). Comparative assessment of handedness for a coordinated bimanual task in chimpanzees (Pan troglodytes), gorillas (Gorilla gorilla) and orangutans (Pongo pygmaeus). Journal of Comparative Psychology, 117(3), 302.

Huber, K. B., & Marsolek, C. J. (2022). Do cerebral motivational asymmetries mediate the relationship between handedness and personality ?. Laterality, 27(1), 21–56.

Karanth, K. U., Srivathsa, A., Vasudev, D., Puri, M., Parameshwaran, R., & Kumar, N. S. (2017). Spatio-temporal interactions facilitate large carnivore sympatry across a resource gradient. Proceedings of the Royal Society B: Biological Sciences, 284(1848), 20161860.

Kashyap, D. K., Tiwari, S. K., & Girl, D. K. (2011). Management of Electrocution in a Langur (Semnopithecus entellus). Intas Polivet, 12(2).

King, J. E., & Landau, V. I. (1993). Reaching in squirrel monkeys. Primate laterality: Current behavioral evidence of primate asymmetries, 107.

MacNeilage, P. F. (1998). Towards a unified view of cerebral hemispheric specializations in vertebrates. In Comparative neuropsychology (ed. A. D. Milner), pp. 167–183. Oxford, UK: Oxford University Press.

MacNeilage, P. F., Studdert-Kennedy, M. G., & Lindblom, B. (1987). Primate handedness reconsidered. Behavioral and Brain Sciences, 10(2), 247–263.

Meguerditchian, A., Donnot, J., Molesti, S., Francioly, R., & Vauclair, J. (2012). Sex difference in squirrel monkeys’ handedness for unimanual and bimanual coordinated tasks. Animal Behaviour, 83(3), 635–643.

Meguerditchian, A., Vauclair, J., & Hopkins, W. D. (2013). On the origins of human handedness and language: a comparative review of hand preferences for bimanual coordinated actions and gestural communication in nonhuman primates. Developmental Psychobiology, 55(6), 637–650.

Mittra, E. S., Fuentes, A., & McGrew, W. C. (1997). Lack of hand preference in wild Hanuman langurs (Presbytis entellus). American Journal of Physical Anthropology: The Official Publication of the American Association of Physical Anthropologists, 103(4), 455–461.

Narasimmarajan, K., Puia, L., & Barman, B. B. (2012). Population density, Group size and Abundance of Hanuman langurs (Semnopithecus entellus) in Melghat Tiger Reserve, Maharashtra, Central India. NeBIO, 3(1), 84–88.

Patel, B. A. (2010). Functional morphology of cercopithecoid primate metacarpals. Journal of human evolution, 58(4), 320–337.

Patil, S., & Modse, S. (2018). Food and feeding in hanuman langurs (Semnopithecus entellus) of bidar District Karnataka. Journal of Progressive Agriculture, 9(2), 6–17.

Prieur, J., Pika, S., Barbu, S., & Blois-Heulin, C. (2016). Gorillas are right-handed for their most frequent intraspecific gestures. Animal Behaviour, 118, 165–170.

Rahman, M. M., Jaman, M. F., Khatun, M. T., Alam, S. M. I., Kayum, A. R. M. R., & Uddin, M. (2015). Substrate utilization by Bengal Sacred langur Semnopithecus entellus (Dufresne, 1797) in Jessore, Bangladesh: effect of resource type on feeding in urban and rural groups. Int. J. Pure Appl. Zool, 3, 162–172.

Regaiolli, B., Spiezio, C., & Hopkins, W. D. (2016). Hand preference on unimanual and bimanual tasks in strepsirrhines: The case of the ring□tailed lemur (Lemur catta). American journal of primatology, 78(8), 851–860.

Rogers, L. J. & Andrew, R. J. (2002). Comparative vertebrate lateralization. Cambridge University Press.

Rogers, L. J. (2002). Lateralised brain function in anurans: Comparison to lateralisation in other vertebrates. Laterality: Asymmetries of Body, Brain and Cognition, 7(3), 219–239.

Rogers, L. J. (2009). Hand and paw preferences in relation to the lateralized brain. Philosophical Transactions of the Royal Society B: Biological Sciences, 364(1519), 943–954.

Sayers, K., & Norconk, M. A. (2008). Himalayan Semnopithecus entellus at Langtang National Park, Nepal: Diet, Activity Patterns, and Resources. International Journal of Primatology, 29(2), 509–530. doi: 10.1007/s10764-008-9245-x

Schweitzer, C., Bec, P., & Blois□Heulin, C. (2007). Does the complexity of the task influence manual laterality in De Brazza’s monkeys (Cercopithecus neglectus)?. Ethology, 113(10), 983–994.

Schweitzer, C., Bec, P., & Blois□Heulin, C. (2007). Does the complexity of the task influence manual laterality in De Brazza’s monkeys (Cercopithecus neglectus) ?. Ethology, 113(10), 983–994.

Spinozzi, G. (2007). Factors affecting manual laterality in tufted capuchins (Cebus apella). Special Topics in Primatology, 5, 204–226.

Spinozzi, G., Lagana, T., & Truppa, V. (2007). Hand use by tufted capuchins (Cebus apella) to extract a small food item from a tube: digit movements, hand preference, and performance. American Journal of Primatology: Official Journal of the American Society of Primatologists, 69(3), 336–352.

Sushma, H. S., & Singh, M. (2006). Resource partitioning and interspecific interactions among sympatric rain forest arboreal mammals of the Western Ghats, India. Behavioral Ecology, 17(3), 479–490.

Vallortigara, G., & Rogers, L. (2005). Survival with an asymmetrical brain: advantages and disadvantages of cerebral lateralization. Behavioral and brain sciences.

Westergaard, G. C, & Suomi, S. J. (1996). Hand preference for a bimanual task in tufted capuchins (Cebus apella) and rhesus macaques (Macaco mulatto). Journal of Comparative Psychology, 110, 406–411.

Westergaard, G. C., Champoux, M., & Suomi, S. J. (1997). Hand preference in infant rhesus macaques (Macaca mulatta). Child Development, 68(3), 387–393.

Zhao, D., Gao, X., & Li, B. (2010). Hand preference for spontaneously unimanual and bimanual coordinated tasks in wild Sichuan snub-nosed monkeys: implication for hemispheric specialization. Behavioural Brain Research, 208(1), 85–89.

Zhao, D., Hopkins, W. D., & Li, B. (2012). Handedness in nature: first evidence on manual laterality on bimanual coordinated tube task in wild primates. American Journal of Physical Anthropology, 148(1), 36–44.

